# Lung epithelial CYP1 activity regulates aryl hydrocarbon receptor dependent allergic airway inflammation

**DOI:** 10.1101/2021.09.20.461064

**Authors:** Francesca Alessandrini, Renske de Jong, Maria Wimmer, Ann-Marie Maier, Isis Fernandez, Miriam Hils, Jeroen T. Buters, Tilo Biedermann, Ulrich Zissler, Christian Hoffmann, Julia Esser-von-Bieren, Carsten B. Schmidt-Weber, Caspar Ohnmacht

**Author notes:** contributed equally (Francesca Alessandrini, Renske de Jong). **Correspondence**, Caspar Ohnmacht, Center of Allergy and Environment (ZAUM), Technical University and Helmholtz Center Munich, building 57, room 105, Ingolstaedter Landstrasse 1, D-85764 Neuherberg, Germany.

## Abstract

The lung epithelial barrier serves as a guardian towards environmental insults and responds to allergen encounter with a cascade of immune reactions that can possibly lead to inflammation. Whether the environmental sensor aryl hydrocarbon receptor (AhR) together with its downstream targets cytochrome P450 (CYP1) family members contribute to the regulation of allergic airway inflammation remains unexplored. By employing knockout mice for AhR and for single CYP1 family members, we found that AhR^-/-^ and CYP1B1^-/-^ but not CYP1A1^-/-^ or CYP1A2^-/-^ animals display enhanced allergic airway inflammation compared to WT. Expression analysis, immunofluorescence staining of murine and human lung sections and bone marrow chimeras suggest an important role of CYP1B1 in non-hematopoietic lung epithelial cells to prevent exacerbation of allergic airway inflammation. Transcriptional analysis of murine and human lung epithelial cells indicates a functional link of AhR to barrier protection/inflammatory mediator signaling upon allergen challenge. In contrast, CYP1B1 deficiency leads to enhanced expression and activity of CYP1A1 in lung epithelial cells and to an increased availability of the AhR ligand kynurenic acid following allergen challenge. Thus, differential CYP1 family member expression and signaling via the AhR in epithelial cells represents an immunoregulatory layer protecting the lung from exacerbation of allergic airway inflammation.

## Introduction

The aryl hydrocarbon receptor (AhR) is a ligand-activated transcription factor used by immune cells and non-hematopoietic cells to sense the environment and adapt to physiological processes particularly at and within barrier sites such as the skin, the gut or the lung^1^. Upon ligand binding, cytoplasmic AhR translocates to the nucleus to regulate gene transcription either alone or in a complex with other transcription factors. The AhR thereby acts as an intracellular sensor, which is able to respond to a variety of organic compounds including xenobiotics and tryptophane metabolites both from the host and from microbial/ dietary sources. Interestingly, knockout of the AhR or deprivation of AhR ligands has been shown to alter the proper function of the immune system, particularly intestinal type 3 immunity including Th17 cells, innate lymphoid cells type (ILC)3 and a subset of intraepithelial cells^2–5^.

In the context of allergic diseases, ablation of the AhR has been shown to increase the severity of allergic inflammation, indicating a rather protective role of the AhR in these settings^6–11^. In fact, the AhR agonist Tranilast has been successfully employed as anti-allergic drug^12–14^. Noteworthy, the AhR also regulates ILC2 activity^15^, immunogenicity of dendritic cells^16^, macrophage polarization towards a M2 phenotype via mesenchymal cells^17^ and class-switch recombination as well as IL-10 expression in B cells^18, 19^. Mechanistically, activation of the AhR typically results in upregulation of the cytochrome P450 (CYP1) members *cyp1a1*, *cyp1a2* and *cyp1b1^1^*. CYP enzymes are monooxygenases that catalyze various reactions involved in drug metabolism, synthesis of cholesterol, steroids and other lipids. Importantly, a polymorphism in the *CYP1B1* gene has been linked to enhanced risk of bronchial asthma^20^. However, whether and how AhR activity and particularly individual cytochrome P450 members play a role in allergic airway inflammation (AAI) has not been addressed so far.

Therefore, we investigated whether knockout of AhR and individual CYP1 family members differentially impact AAI in murine model systems using clinically-relevant, naturally occurring aeroallergens. We show that AhR and CYP1B1 but not CYP1A1 or CYP1A2 deficiency leads to disease exacerbation in preclinical models of AAI. We further show that airway epithelial cells (ECs) express high levels of CYP1B1 and CY1B1 expression in non-hematopoietic cells is required for protection from exacerbated AAI. AhR deficiency results in alteration of gene expression profiles in primary airway ECs at steady state and even more following AAI, but CYP1B1 deficiency did not. We identified enhanced CYP1A1 activity in CYP1B1^-/-^ ECs and altered tryptophane (Trp) metabolite levels as a possible mechanism responsible for the AAI-potentiating effects observed in CYP1B1^-/-^animals. This work reveals a novel protective role of AhR signaling in airway ECs, which may have the potential to serve as a novel target for therapy of allergic asthma.

## Results

### AhR and CYP1B1 deficiency aggravates ragweed-induced allergic airway inflammation

To assess the role of AhR signaling and cytochrome P450 enzymes in AAI, we sensitized AhR^-/-^, CYP1A1^-/-^, CYP1A2^-/-^ and CYP1B1^-/-^ mice to ragweed extract as depicted in Fig 1a. Although the overall detection of serum immunoglobulins was very modest, CYP1B1^-/-^ animals showed a significant increase at the endpoint in total IgE compared to WT (Fig 1b; left panel) and Amb a 1-specific IgG1 level compared to PBS, WT and CYP1A2^-/-^ (Fig 1b; right panel). A tendency towards increased IgE and Amb a 1-specific IgG1 levels was observed also for AhR^-/-^ mice albeit not reaching significance (Fig 1b). Enhanced BAL cell infiltration could be observed in AhR^-/-^ and CYP1B1^-/-^ compared to PBS and WT but not to CYP1A1^-/-^ and CYP1A2^-/-^ animals (Fig 1c). Differential cell counts revealed that this effect was predominantly due to enhanced infiltration of eosinophils, lymphocytes and macrophages into the bronchoalveolar space (Fig 1c). Additionally, CYP1B1^-/-^ animals revealed enhanced frequencies of Gata3^+^Foxp3^−^ T helper (Th2) cells in lung tissue compared to PBS, WT, AhR^-/-^ and CYP1A1^-/-^ (Fig 1d). Gene expression analysis of key Th1/Th2-associated cytokines and CCL11 in lung tissue revealed a tendency for increased IL-4 expression in AhR^-/-^ and CYP1B1^-/-^ animals, but overall, no significant differences were detected between the investigated genotypes (Fig 1e). Histological analysis of lung tissue showed strongest inflammatory infiltration and mucus hypersecretion in AhR^-/-^ - and CYP1B1^-/-^ animals, although significantly only compared to PBS and not to WT or to other genotypes (Fig 1f). Altogether, these results demonstrate that AhR^-/-^ and CYP1B1^-/-^ animals suffer from a stronger AAI upon chronic exposure to ragweed extract compared to WT and to the other investigated genotypes.

**Figure 1.**
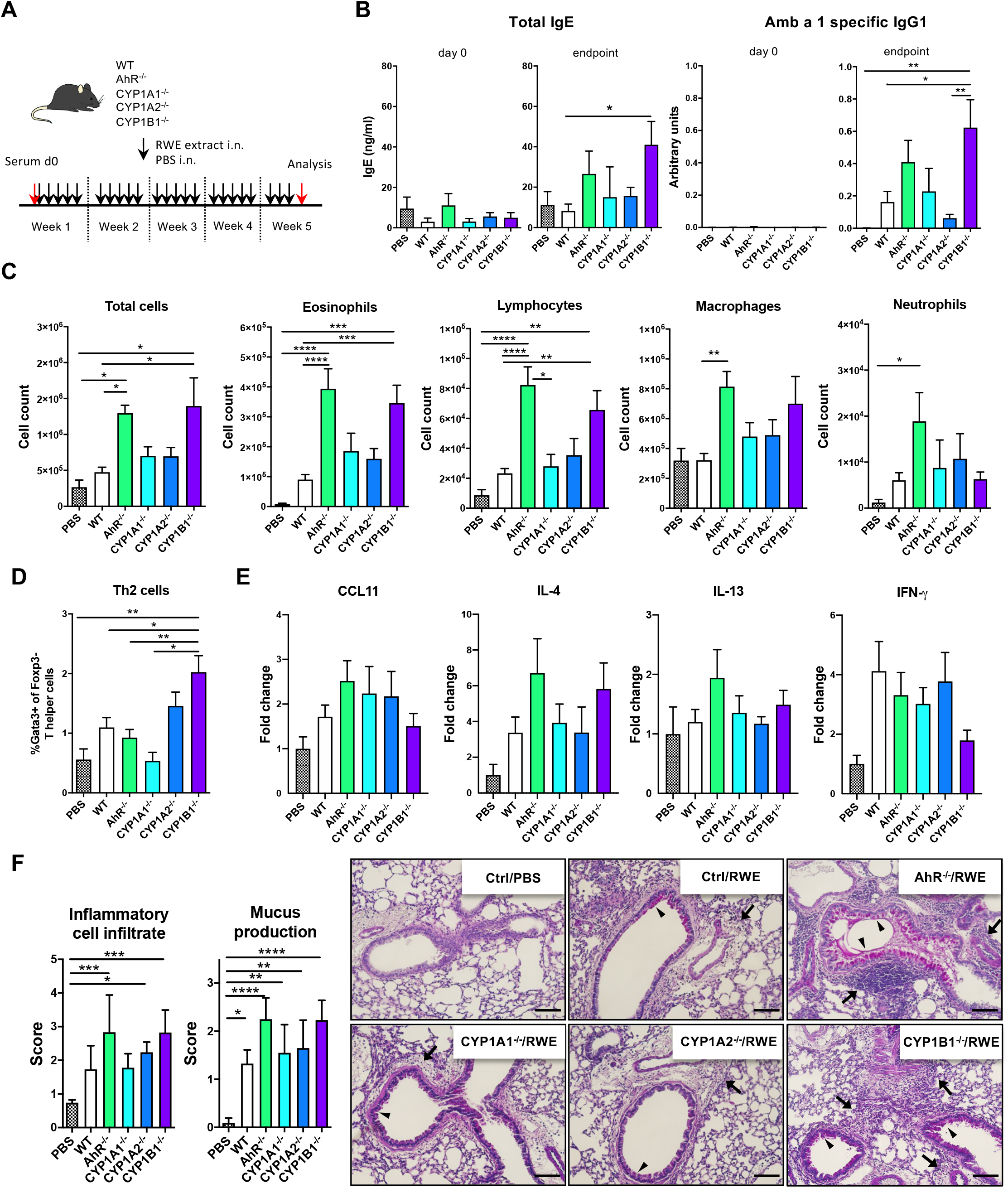
AhR and CYP1B1 deficiency aggravates pollen-induced allergic airway inflammation. **(A)** Experimental setup. Mice of indicated genotypes were exposed for five weeks to ragweed extract (RWE). Wildtype mice exposed to PBS served as negative controls (PBS). **(B)**Total IgE (left) at day 0 and endpoint and Amb a 1-specific IgG1 (right), respectively. Mean ± SEM Total IgE (PBS n=10; WT: n=20; AhR^-/-^ n=17; CYP1A1^-/-^ n=5; CYP1A2^-/-^ n=5; CYP1B1^-/-^ n=20 from up to 3 independent experiments). Amb a 1-specific IgG1 (PBS n=10; WT n=17; AhR^-/-^ n=11; CYP1A1^-/-^ n=6; CYP1A2^-/-^ n=6; CYP1B1^-/-^ n=13 from up to 3 independent experiments). **(C)** Total and differential BAL cell counts. Mean ± SEM (PBS n=9; WT n= 21; AhR^-/-^ n=15; CYP1A1^-/-^ n=6; CYP1A2^-/-^ n=6; CYP1B1^-/-^ n=15 from up to 3 independent experiments). **(D)** Th2 cell frequency. Mean ± SEM (PBS n=6; WT n= 18; AhR^-/-^ n=12; CYP1A1^-/-^ n=4; CYP1A2^-/-^ n=4; CYP1B1^-/-^ n=14 from up to 3 independent experiments). **(E)** CCL11, IL-4, IL-13 and IFN-g expression in whole lung tissue. Mean ± SEM (PBS n=5; WT n= 18; AhR^-/-^ n=16; CYP1A1^-/-^ n=6; CYP1A2^-/-^ n=6; CYP1B1^-/-^ n=19 from up to 3 independent experiments). **(F)** Histological scores (left) and representative lung sections (right) Arrows: inflammatory infiltrate; arrowheads: mucus hypersecretion; scale bar: 100μm. Mean ± SD (n=5 mice/group). **(B-F)** One-way ANOVA with Tukey’s multiple comparisons test. *p < 0.05, **p < 0.01, ***p < 0.001, ****p < 0.0001

### AhR and CYP1B1 deficiency aggravates house dust mite-induced allergic airway inflammation

In order to assess the allergen specificity of our finding in the ragweed model, we next exposed WT, AhR^-/-^ and CYP1B1^-/-^ animals to HDM-induced AAI (Fig 2a). Here, a more pronounced increase of total serum IgE and a significant increase of Der f specific IgG1 was found both in AhR^-/-^ and CYP1B1^-/-^ compared to WT mice (Fig 2b, 2g). Further, we found enhanced lung cellular infiltration in the BALF of AhR^-/-^ and CYP1B1^-/-^ animals predominantly due to eosinophils and lymphocytes and macrophages (Fig 2c, 2h). No difference in BALF neutrophils was detected (data not shown). In contrast, increased frequencies of Th2 cells could be observed in lung tissue of both genotypes compared to WT (Fig 2d, 2i). AhR^-/-^ mice showed three- to fourfold higher levels of IL-4 and IL-13 as well as of CCL11 in lung tissue and a slight increase of IL-17a, whereas CYP1B1^-/-^ mice showed only slightly elevated expression of IL-4, a strong increase of IL-17a and no variations in CCL11 and IL-13 compared to WT (Fig. 2e and 2j). IFN-γ levels were not affected in either genotype (data not shown). Lastly, histological analysis of HDM-exposed lungs revealed enhanced peribronchiolar and perivascular infiltration and mucus hypersecretion in both AhR^-/-^ and CYP1B1^-/-^ animals compared to WT (Fig 2f and 2k). Altogether our *in vivo* data from two different adjuvant-free AAI models indicate that both AhR and its downstream target CYP1B1 are essential to prevent excessive AAI in a non-allergen-specific manner.

**Figure 2.**
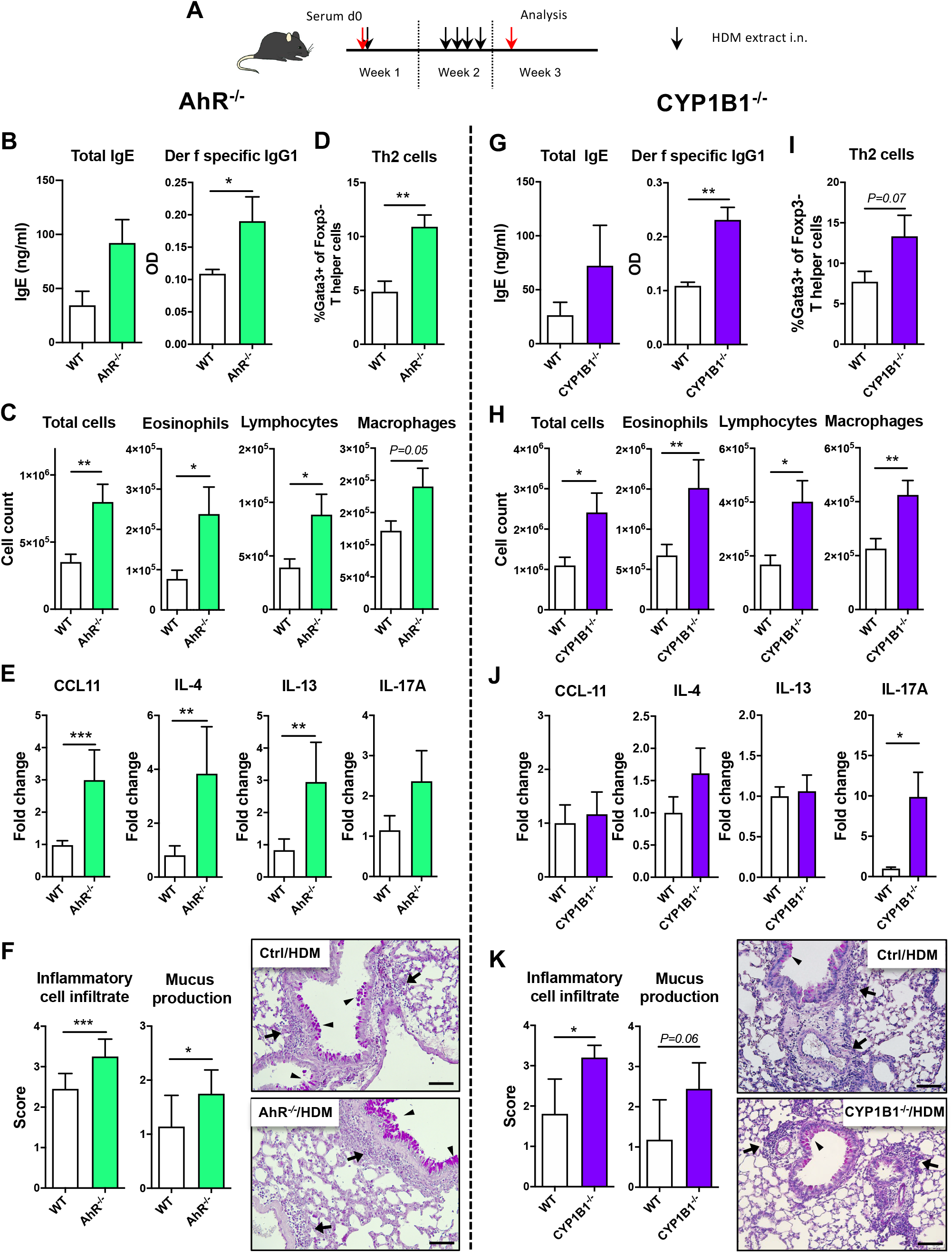
AhR and CYP1B1 deficiency aggravates HDM-induced allergic airway inflammation. **(A)** Experimental setup. AhR^-/-^ (left) and CYP1B1^-/-^ (right) mice were exposed to house dust mite (HDM) extract as indicated. No PBS-treated groups were included because neither WT, AhR^-/-^ nor CYP1B1^-/-^ show signs of airway inflammation after PBS treatment only. **(B** and **G)** Total IgE (left) and *Der f*-specific IgG1 (right) at endpoint. Mean ± SEM. AhR^-/-^ vs WT: total IgE n=6/group; Der f sp. IgG1 (WT n=8, AhR^-/-^ n=7); CYP1B1^-/-^ vs WT: total IgE n=10/group; Der f sp. IgG1 (WT n=8, CYP1B1^-/-^ n=7), from two independent experiments. **(C** and **H)** Total and differential BAL cell counts. Mean ± SEM. AhR^-/-^ vs WT: n=10/group from two independent experiments. CYP1B1^-/-^ vs WT: n=10/group from two independent experiments. **(D** and **I)** Th2 cell frequency. Mean ± SEM. AhR^-/-^ vs WT: n=5/group; CYP1B1^-/-^ vs WT: n=10/group from two independent experiments. **(E** and **J)** CCL11, IL-4, IL-13, IL-17a expression in whole lung tissue. Mean ± SEM. AhR^-/-^ vs WT: n=6-8/group. CYP1B1^-/-^ vs WT: n=6-10/group; pooled from up to two independent experiments. **(F** and **K)** Histological scores (left) and representative lung sections (right) at endpoint in AhR^-/-^ and CYP1B1^-/-^, respectively. Arrows: inflammatory infiltrate; arrowheads: mucus hypersecretion; scale bar: 100μm. Mean ± SD. AhR^-/-^ vs WT: n=8-11/group from two independent experiments. CYP1B1^-/-^ vs WT: n=4-5/group from two independent experiments. **(B-K)** Student’s t-test (unpaired, two-tailed). *p < 0.05, **p < 0.01, ***p < 0.001

### CYP1B1 is primarily expressed in airway epithelial cells

Given the selective effect of CYP1B1expression on the allergic inflammatory response, we then aimed to identify the cell type(s) that primarily express CYP1B1 in the lung and to evaluate if our results can be translated to human. Therefore, we first separated hematopoietic (CD45^+^) and non-hematopoietic (CD45^−^ cells) from the lungs of naïve WT, AhR^-/-^ and CYP1B1^-/-^ animals. Strikingly, non-hematopoietic WT lung cells expressed roughly 100-200 times higher levels of CYP1B1 compared to hematopoietic cells at baseline (Fig 3a). Interestingly, CYP1B1 expression was not affected by the absence of AhR (Fig 3a). Immunofluorescence staining of murine lung tissue showed that CYP1B1 stained within bronchial epithelial cells (EC), similarly in control and allergic animals (Fig 3b). Also, in human lung sections, CYP1B1 stained bronchial ECs (Fig 3c). In contrast, CYP1B1 did not stain alveolar ECs in mouse or human lung sections (Fig 3b, c). These data are in line with single cell expression data demonstrating CYP1B1 but not CYP1A1 or CYP1A2 expression across different subsets of human lung epithelial and fibroblastic cells (Supplementary Fig S1).

**Figure 3.**
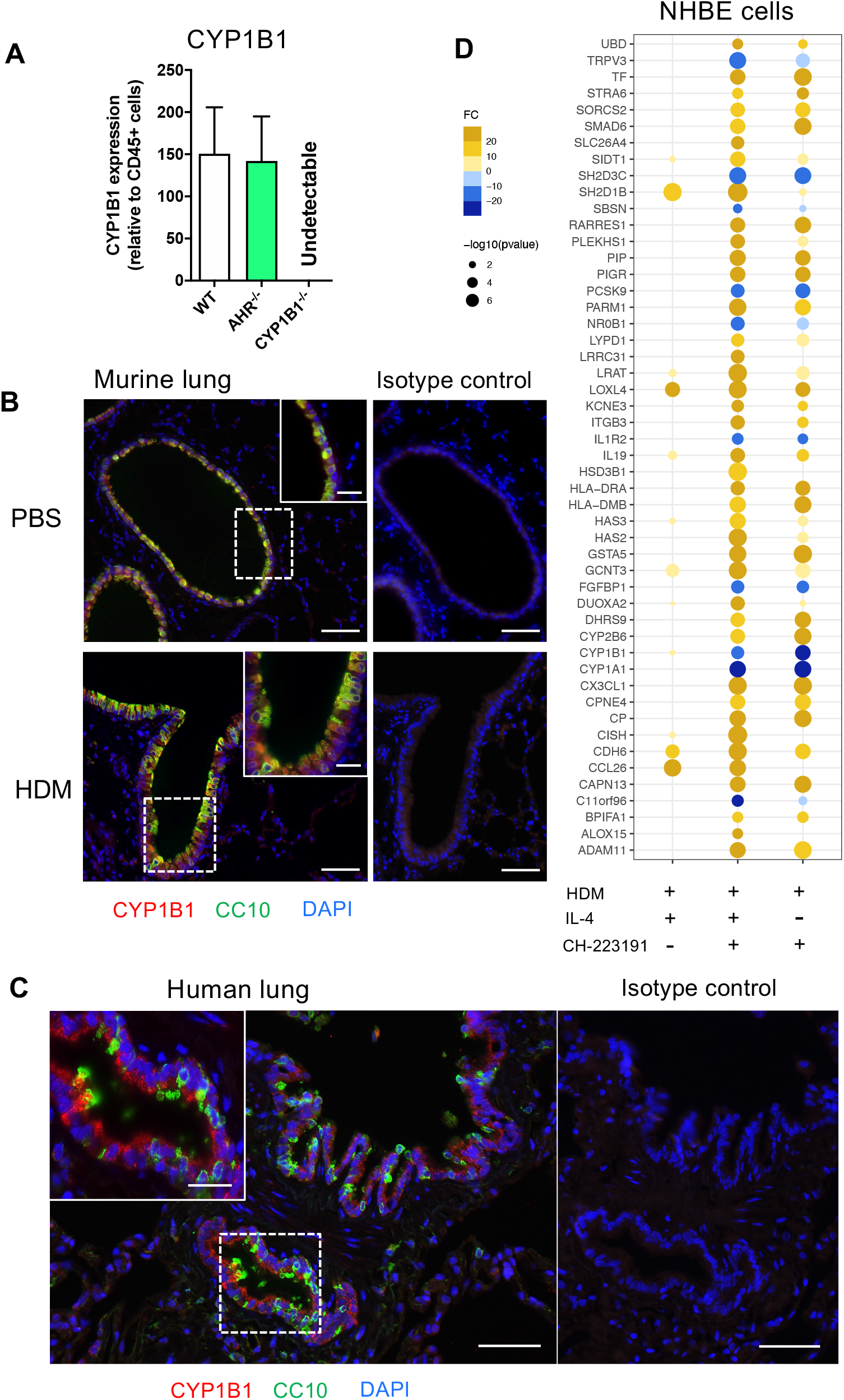
Non-hematopoietic lung cells express CYP1B1. **(A)** CYP1B1 expression in CD45^−^ relative to CD45^+^ lung cells from WT, AhR^-/-^ and CYP1B1^-/-^ mice; Mean±SEM. WT n=6, AhR and CYP1B1^-/-^ n=2. **(B)** Immune fluorescence staining of PBS- and HDM-challenged WT murine lungs or **(C)**human lung sections. Red: CYP1B1, Green: CC10, Blue: DAPI. Scale bar: 50μm or 20μm (insets). (**D**) NHBEs were stimulated for 24h with HDM extract in the presence of the AhR inhibitor CH-223191, IL-4 or a combination of both. Results display top 25 upregulated and top 10 downregulated DEGs identified in the IL-4- and CH-223191-treated group relative to HDM only (center). These DEGs are also shown in NHBEs treated solely with IL-4 (left) or CH-223191 (right), respectively. Mean fold change of 4 independent donors. NHBE, normal human bronchial epithelial cell.

To assess whether human lung ECs are directly regulated by AhR signaling, normal human bronchial epithelial cells (NHBE) were stimulated for 24 h with HDM in the presence of IL-4 (to mimic an ‘allergic’ status), an AhR inhibitor or the combination of both. As expected, microarray analysis revealed a strong effect of HDM exposure compared to non-exposed cells (not shown). Whereas IL-4 strongly stimulated the expression of typical pro-allergic genes such as *CCL26* or *IL13R2* and interestingly also *AHR* (Fig 3d and Supplementary Table S3), AhR inhibition led to a more drastic change in differentially expressed genes (DEGs) compared to IL-4 (Supplementary Table S3). We then focused the analysis on the DEGs mostly altered solely by the inhibition of AhR during HDM exposure. As expected, inhibition of AhR led to downregulation of the AhR target genes *CYP1A1* and *CYP1B1* irrespective of whether IL-4 was present or not. Noteworthy, the inhibition of AhR lead to a strong upregulation of genes critical for allergic tissue inflammation, e.g. cell recruitment (*CX3CL1*), barrier integrity/protection (*ITGB3*, *PIGR*), antigen presentation (*HLA-DRA*, *HLA-DMB*), Th2 differentiation (*IL19*) and lipid mediator synthesis (*ALOX15*) (Fig 3d and Supplementary Table S3).

Thus, bronchial ECs seem to be the major CYP1B1-expressing cell type in murine and human lung. Furthermore, AhR activity in allergen-exposed NHBEs regulates a number of genes associated to AAI.

### Non-hematopoietic expression of CYP1B1 protects from exacerbation of HDM-induced allergic airway inflammation

Given the predominant expression of CYP1B1 in mouse and human bronchial ECs, we then investigated whether non-hematopoietic expression of CYP1B1 is sufficient to prevent exacerbation of AAI. To this end, we generated CYP1B1^-/-^ and WT bone marrow chimeras (Fig 4a) and sensitized them to HDM, as depicted in Fig. 2a. Strikingly, only CYP1B1^-/-^ chimeras with WT hematopoietic cells mirrored previously detected parameters of elevated AAI in full CYP1B1^-/-^ mice, including serum immunoglobulin response (Fig 4b), BAL total cell numbers, particularly eosinophils and lymphocytes (Fig 4c), lung-resident Th2 cells (Fig 4d), IL-4 and IL-17 in BALF (Fig 4e) and histological scores in lung sections (Fig 4f). Conversely, WT chimeras with CYP1B1^-/-^ hematopoietic cells showed much milder symptoms of AAI. Thus, CYP1B1 expression in non-hematopoietic cells is primarily responsible to prevent exaggerated AAI after exposure to HDM.

**Figure 4.**
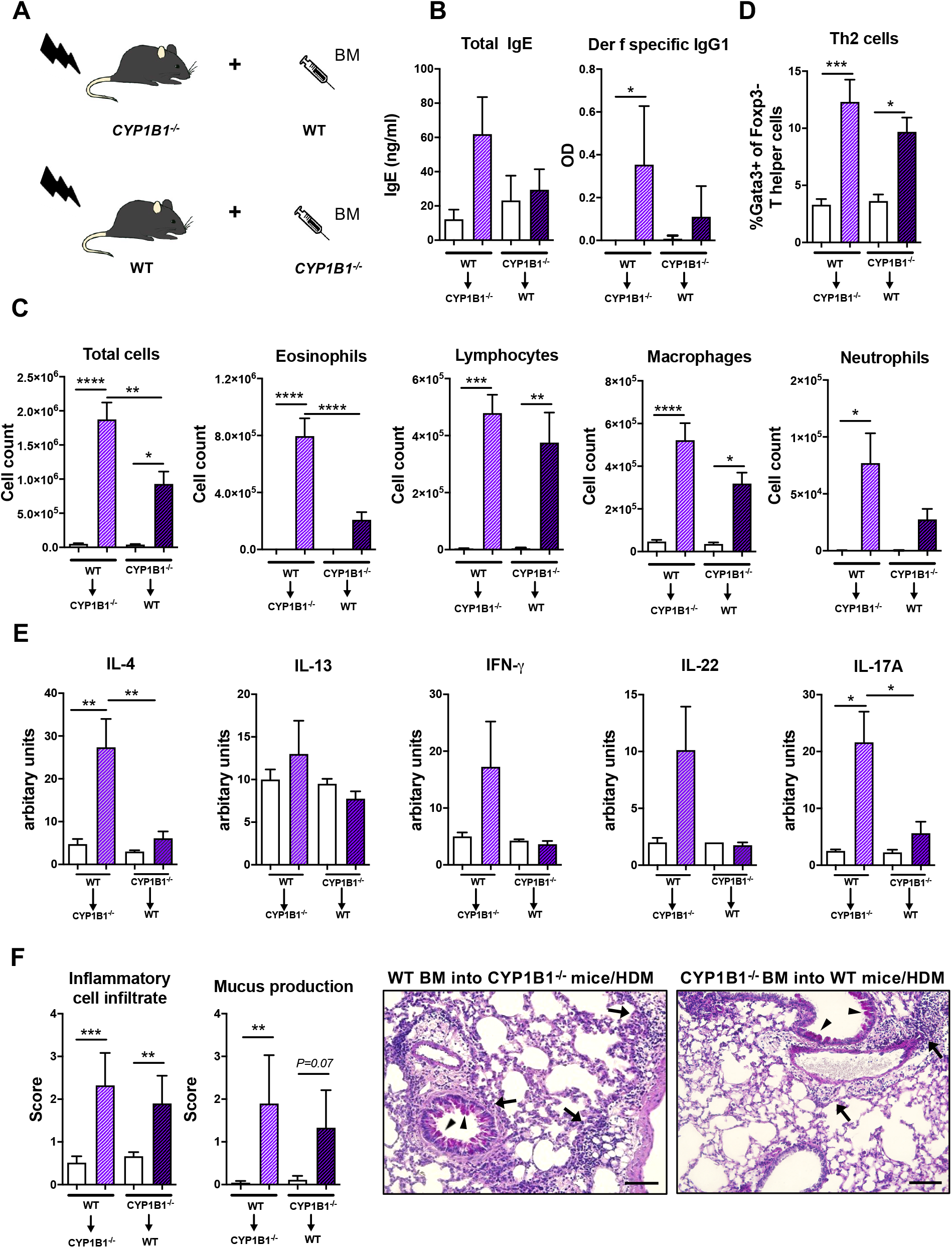
Non-hematopoietic CYP1B1 expression prevents exacerbation of HDM-induced allergic airway inflammation. **(A)** Schematic representation of the generation of WT and CYP1B1^-/-^ bone marrow chimeras, which were subsequently treated with HDM or PBS (for setup see Fig 2a). **(B)** Total IgE (left) and *Der f*-specific IgG1 (right) at endpoint. IgE: Mean ± SEM (WT-BM in CYP1B1^-/-^ PBS n=9; WT-BM in CYP1B1^-/-^ HDM n=13; CYP1B1^-/-^-BM in WT PBS n=8; CYP1B1^-/-^-BM in WT HDM n=11) from 2 independent experiments. *Der f*-specific IgG1: Mean ± SD, n=5/group. **(C)** Total and differential BAL cell counts; Mean ± SEM (WT-BM in CYP1B1^-/-^ PBS n=9; WT-BM in CYP1B1^-/-^ HDM n=13; CYP1B1^-/-^-BM in WT PBS n=8; CYP1B1^-/-^-BM in WT HDM n=11) from 2 independent experiments. **(D)** Th2 cell frequency. Mean ± SEM (WT-BM in CYP1B1^-/-^ PBS n=9; WT-BM in CYP1B1^-/-^ HDM n=13; CYP1B1^-/-^-BM in WT PBS n=8; CYP1B1^-/-^-BM in WT HDM n=11) from 2 independent experiments. **(E)** IL-4, IL-13, IFN-g, IL-22 and IL-17A protein levels in whole lung tissue. Mean ± SEM (WT-BM in CYP1B1^-/-^ PBS n=4; WT-BM in CYP1B1^-/-^ HDM n=8; CYP1B1^-/-^-BM in WT PBS n=4; CYP1B1^-/-^-BM in WT HDM n=8) from up to 2 independent experiments. **(F)** Histological scores (left) and representative lung sections (right) Arrows: inflammatory infiltrate; arrowheads: mucus hypersecretion; scale bar: 100μm. Mean ± SD (WT-BM in CYP1B1^-/-^ PBS n=5; WT-BM in CYP1B1^-/-^ HDM n=4; CYP1B1^-/-^-BM in WT PBS n=5; CYP1B1^-/-^-BM in WT HDM n=4). **(B-F)** ANOVA with Tukey’s multiple comparisons. BM, bone marrow. *p < 0.05, **p < 0.01, ***p < 0.001, ****p < 0.0001

### Transcriptional comparison indicates AhR-but not CYP1B1-dependent regulation of bronchial epithelial cells

To get a deeper insight into the molecular mechanism responsible for AhR- and CYP1B1-dependent effects, we performed RNA sequencing of sort-purified primary lung ECs (CD45^−^CD31^−^EpCAM^+^live^+^) at steady state and after HDM-elicited-AAI. The analysis of the number of DEGs in AhR^-/-^ and CYP1B1^-/-^ relatively to WT showed a stronger impact of the deficiency of AhR compared to CYP1B1 in gene regulation, especially following HDM exposure (Fig 5a). In fact, at steady state, we detected only few DEGs between WT and AhR-deficient ECs. Interestingly, some of these genes have been linked to regulation of circadian clock (*Nr1d1*, *Nr1d2*, *Cry1*) (Fig 5b). Following HDM-induced allergy, ECs from AhR^-/-^ showed a strongly altered gene expression profile, contrarily to CYP1B1^-/-^, which showed only low DEGs compared to WT with and none without allergy (Fig 5b and Supplementary Table S4). KEGG pathway analysis revealed an overall upregulation of genes involved in cell-to-cell contact and TGF-β signaling whereas genes associated to pattern recognition (TLR-like and NOD-like receptor signaling), chemokine receptor signaling and JAK-STAT signaling pathway were overall downregulated (Fig 5c and Supplementary Table S5). Overall, these data indicate that AhR, but not CYP1B1, is a major transcriptional regulator of ECs function, particularly after exposure to aeroallergens.

**Figure 5.**
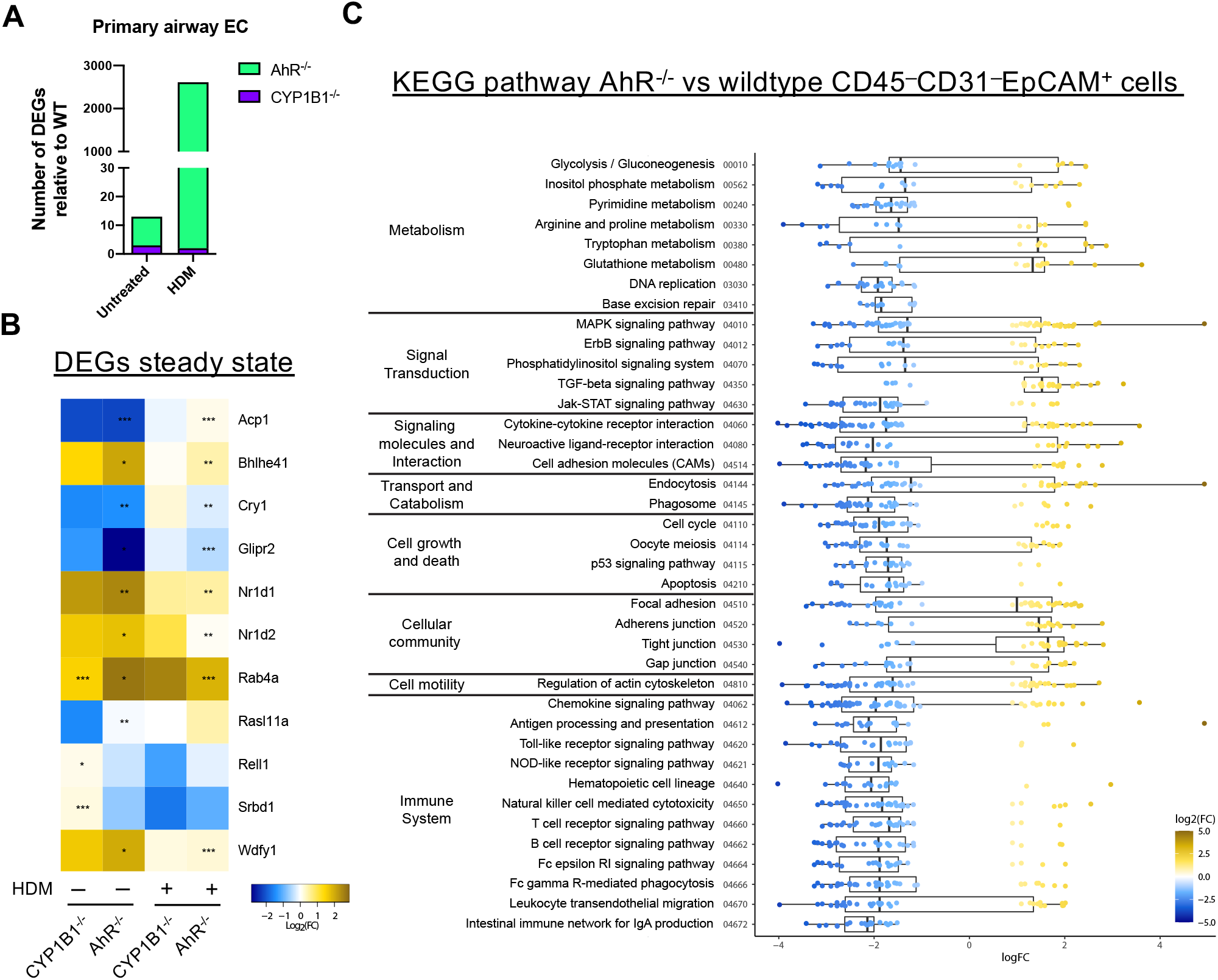
AhR regulates gene expression in airway epithelial cells of sensitized mice. **(A)** Total DEGs in AhR^-/-^ and CYP1B1^-/-^ epithelial cells (EC) relatively to WT from untreated or allergic (HDM) mice. (**B** and **C**) RNAseq analysis of sort-purified EC (CD45^−^CD31^−^EpCAM^+^live^+^) of AhR^-/-^ and CYP1B1^-/-^ mice. **(B)** Heatmap of all genes differently expressed in AhR^-/-^ or CYP1B1^-/-^ versus WT EC at steady state and their expression in HDM-exposed animals. **(C)** KEGG pathway analysis of DEGs between AhR and WT EC of HDM-exposed animals. Only pathways with at least 5 DEGs are shown.

### Absence of CYP1B1 leads to enhanced transcription and activity of CYP1A1 and to altered tryptophane metabolites levels

Our results indicating a non-transcriptional effect of CYP1B1 deficiency in ECs prompted us to investigate underlying mechanisms responsible for the enhanced allergic phenotype in CYP1B1^-/-^ mice. Since artificial overexpression of CYP1A1 is known to limit availability of AhR agonists *in vivo*^5^, we measured the expression of CYP1A1 in the absence of CYP1B1. Interestingly, lung tissue of CYP1B1^-/-^ animals showed higher CYP1A1 expression levels compared to WT both at steady state and in tendency also following RWE-induced AAI, whereas CYP1A1 was not detected in AhR^-/-^ or CYP1A1^-/-^ lung tissue (Fig 6a, b). Similarly, higher CYP1A1 expression compared to WT could be observed in CYP1B1^-/-^ lung tissue following HDM-induced AAI (Fig 6c). Notably, this effect was even more pronounced in purified non-hematopoietic (CD45^−^) cells from HDM-allergic lungs with a roughly 20-fold higher CYP1A1 expression in the absence of CYP1B1 (Fig. 6d).

**Figure 6.**
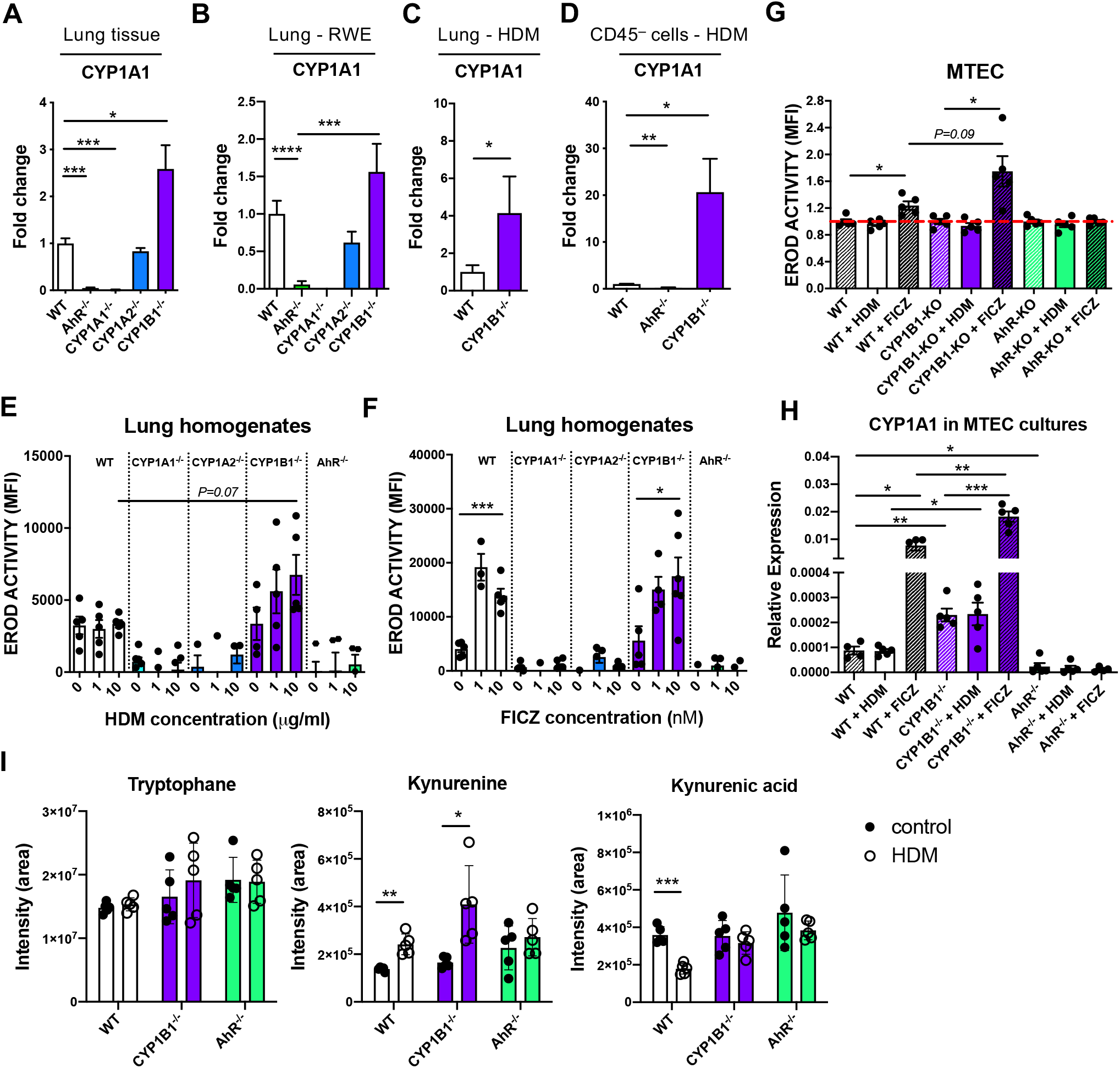
Functional consequences of CYP1B1 deficiency in airway epithelial cells. **(A)** CYP1A1 expression in total lung untreated tissue. Mean ± SEM (WT n=6, AhR^-/-^ n=4, CYP1A1^-/-^ n=2, CYP1A2^-/-^ n=2, CYP1B1^-/-^ n=6). **(B)** CYP1A1 expression in total lung tissue sensitized to RWE. Mean ± SEM (WT n=17, AhR^-/-^ n=18, CYP1A1^-/-^ n=6, CYP1A2^-/-^ n=6, CYP1B1^-/-^ n=20) from up to 3 independent experiments. **(C**) CYP1A1 expression in total lung tissue sensitized to HDM. Mean ± SEM (WT n=10; CYP1B1^-/-^ n=5) from up to 2 independent experiments. **(D)** CYP1A1 expression in CD45^−^ cells isolated from HDM-sensitized of indicated genotypes. Mean ± SEM (WT n=6, AhR^-/-^ n=3, CYP1B1^-/-^ n=7). **(E)** EROD assay in total lung homogenates of indicated genotypes after stimulation with HDM. Mean ± SEM (n/concentration: WT n=5, CYP1A1^-/-^ n=6, CYP1A2^-/-^ n=3, CYP1B1^-/-^ n=5, AhR^-/-^ n=4). **(F)** EROD assay in total lung homogenates of indicated genotypes after stimulation with FICZ. Mean ± SEM (n/concentration: WT n=5, CYP1A1^-/-^ n=6, CYP1A2^-/-^ n=4, CYP1B1^-/-^ n=5, AhR^-/-^ n=4). **(G)** EROD assay of murine MTEC of the indicated genotypes after stimulation with HDM (10μg/mL) or FICZ (10nM). Mean ± SEM; n=5/group/treatment. **(H)** CYP1A1 expression of murine MTEC of the indicated genotypes after stimulation with HDM (10μg/mL) or FICZ (10nM). Mean ± SEM; n=5/group/treatment. **(I)** Targeted LC-MS/MS measurement of tryptophane, kynurenine and kynurenic acid in lung homogenates of the indicated genotype with or without AAI. Mean ± SD; n=5/group/treatment. (**A**-**D)** Student’s t-test (unpaired, two-tailed); **(E-I)** Student’s t-test (unpaired, two-tailed) with Welch’s correction. MTEC, mouse tracheobronchial epithelial cells. *p < 0.05, **p < 0.01, ***p < 0.001, ****p < 0.0001

To assess whether the HDM extract used for sensitization directly activates the AhR in ECs, we stimulated lung homogenates from all investigated genotypes with HDM extract or with the prototypical AhR ligand FICZ and compared the resulting EROD activity. HDM induced a slight, yet non-significant increase of EROD activity only in CYP1B1^-/-^ animals compared to WT (Fig 6e). In contrast, FICZ induced EROD activity indistinctively in WT and CYP1B1^-/-^ but not in all other genotypes (Fig 6f). As CYP1B1 was predominantly expressed in ECs (Fig 3), we measured EROD activity in primary murine tracheobronchial ECs (MTECs). Here, HDM did not elicit EROD activity in any background, whereas FICZ induced highest EROD activity in CYP1B1^-/-^ compared to WT, but not in AhR^-/-^ MTECs (Fig 6g). As expected, baseline CYP1A1 expression was higher in CYP1B1^-/-^ MTEC cultures compared to WT and further increased following FICZ stimulation (Fig 6h). Since the HDM extract used for sensitization lacked AhR activation properties, we investigated whether endogenously produced AhR ligands would be affected by a disrupted AhR-CYP1 axis. Therefore, the endogenous AhR ligands of the Trp pathway kynurenine (Kyn) and kynurenine acid (KynA) ^21, 22^ were measured in lung homogenates at steady state and following AAI. Interestingly, Kyn levels increased in WT and CYP1B1^-/-^ lung cells after AAI while this did not reach significance for AhR^-/-^ (Fig 6i). In contrast, KynA levels decreased after AAI in lungs of WT mice, but not in AhR^-/-^ nor in CYP1B1^-/-^ lungs (Fig 6i). In summary, these results point towards a cross-regulation of different CYP1 family members in bronchial ECs and to the employment of selective endogenous AhR ligands in regulating AhR activity in lung tissue following HDM-induced AAI.

## Discussion

Here, we provide evidence of an important role of the AhR-CYP1 axes in preventing exacerbation of AAI using two adjuvant-free murine models with clinically-relevant aeroallergens. In fact, ablation of the AhR as well as of the immediate AhR response-gene CYP1B1 resulted both in enhanced AAI upon exposure to both RWE and HDM, as characterized by serum immunoglobulins, lung inflammatory cell infiltration and mucus hypersecretion. Our results confirm a recent study showing enhanced AAI and remodeling in an ovalbumin-induced chronic asthma model using AhR^-/-^ compared to WT mice^10^ and complete the picture adding the role of the CYP1 gene family. Whether a similar mechanism is present in humans remains to be elucidated. Still, CYP1B1 expression was clearly confined to bronchial and bronchiolar ECs in mouse and human, in line with early reports showing upregulation of CYP1B1 expression upon exposure to AhR ligands within club cells of peripheral airways^23, 24^. Besides the well-known upregulation of CYP1B1 in an AhR-dependent manner, we also found basal CYP1B1 expression independent of AhR (Fig. 3a) which has also been reported earlier in different settings^25, 26^. Thus, we suggest that CYP1B1 is most likely active in steady state conditions but inflammation may potentiate CYP1B1 (and CYP1A1) expression and activity in an AhR-dependent manner.

Functionally, we demonstrate that a well-balanced AhR-CYP1 activity in non-hematopoietic cells is decisive in the protection from disease exacerbation, as only CYP1B1^-/-^ chimeras with WT hematopoietic cells reproduced the previously detected parameters of elevated AAI in full CYP1B1^-/-^ mice. As opposed to CYP1B1, we did not observe any consistent effect of CYP1A1 nor CYP1A2 deficiency in AAI, conforming to their low expression in bronchial ECs (Fig S1). Noteworthy, previous studies have already suggested a protective role of AhR in skin and intestinal inflammatory disorders^6, 8, 27^. Interestingly, immune regulation in the skin has been associated to AhR activity in non-hematopoietic cells, presumably keratinocytes^6^, confirming the critical role of AhR in epithelium as an interface between environment and mucosal immunity.

### What is the consequence of the paucity of AhR signaling in lung epithelial cells?

Our transcriptional analysis of primary lung epithelial cells from non-allergic mice revealed only relatively few differentially expressed genes in the absence of AhR signaling and almost none in the absence of CYP1B1. Surprisingly, the majority of these AhR-dependent genes indicates a deregulation of the peripheral circadian clock. Interestingly, some key effector cells in allergy, such as mast cells, show circadian regulation patterns^28^ and ablation of circadian clock genes in lung ECs has been shown to aggravate inflammation^29, 30^. Whether disruption of peripheral circadian clock networks has a functional impact in our model should be matter of future investigations. In contrast to the relatively few DEGs found in airway ECs from AhR^-/-^ mice at steady-state, we identified strong differences in gene expression of airway ECs from AhR^-/-^ but not CYP1B1^-/-^ animals following HDM exposure. Pathway analysis of these DEGs revealed a role for AhR to ensure epithelial barrier function, limit TGF-β receptor signaling and regulate a number of pathways associated to pattern recognition receptor or cytokine/chemokine signaling^31, 32^. Some of these pathways may not be regulated directly by the AhR, but are rather a consequence of the enhanced AAI observed in AhR^-/-^ mice. On the other hand, the lack of DEGs in ECs from healthy and allergic CYP1B1^-/-^ animals, despite the enhanced AAI similar to AhR^-/-^, suggests that CYP1B1 deficiency in airway ECs affects hematopoietic cell types by other means. To investigate the mechanism responsible for the key role of CYP1B1 in airway ECs, we first assessed the expression of CYP1A1 in the absence of CYP1B1. As our results indicate enhanced activity of CYP1A1 in lung epithelial cells in the absence of CYP1B1, they point towards a cross regulation of CYP1 family members in lung ECs. This finding is in line with earlier studies showing that lungs and livers of CYP1B1^-/-^ animals stimulated with carcinogenic agents had elevated levels of CYP1A1 expression^33, 34^. As aeroallergens typically interact with the epithelium as particles containing multiple molecules, we also tested whether HDM extracts were able to activate the AhR pathway directly, but did not find functionally relevant quantities of AhR ligands in HDM extracts able to activate WT cells. This suggests that naturally occurring activating ligands of AhR, e.g. of the Trp pathway^35, 36^, may be sufficient for AhR-dependent regulation of AAI. To test this hypothesis, we conducted a targeted metabolomics approach to quantify key endogenous AhR ligands of the Trp pathway in lung tissue of WT compared to AhR- and CYP1B1-deficient mice at steady state and following HDM-driven AAI. Interestingly, in WT and CYP1B1^-/-^ lungs, but not in AhR^-/-^, levels of the AhR ligand Kyn increased after AAI. As AhR expression in lung ECs is directly induced by IL-4 signaling (Supplementary Table S3 and^37, 38^), enhanced AhR activity during AAI, fueled with endogenous ligands, may serve as a feedback loop to protect airway ECs from excessive inflammation^16, 39^.

Moreover, contrarily to Kyn, the levels of the acid form KynA decreased in WT after AAI. A plausible explanation for this observation could be the preferential use of KynA for AhR-dependent regulation of AAI *in vivo*, as KynA has a higher affinity to AhR compared to Kyn^21^. Interestingly, KynA levels remained high in lungs of CYP1B1^-/-^ and AhR^-/-^ mice after AAI. In our study we focused on the two most relevant AhR ligands but further studies should assess the role of additional host-intrinsic AhR ligands. Degradation of AhR ligands by excessive CYP1A1 activity has already been identified previously as an important feedback mechanism to negatively regulate activity and duration of AhR signaling and can therefore functionally mimic AhR deficiency in the intestinal tract^5^. Thus, enhanced CYP1A1 activity in the absence of CYP1B1 may explain a comparable defect of immune regulation in AhR^-/-^ and CYP1B1^-/-^animals upon repetitive contact with aeroallergens. Indeed, the activity of individual CYP1 members are able to differentially modulate the availability of local AhR ligands thereby regulating AhR activity in hematopoietic cells such as ILC2s or macrophages, known players in AAI expressing AhR^15^, in analogy to a similar scenario recently proposed for intestinal ECs and skin keratinocytes^5, 40^.

In summary, this work extends a role of the AhR in the regulation of lung non-hematopoietic cells. Moreover, it suggests that balanced AhR/CYP1 activity within ECs represents an important layer of immune regulation necessary to prevent exacerbation of allergic responses in the lung and offers novel possibilities for tailor-made strategies in immune regulation at barrier sites.

## Methods

### Mice and human samples

The following genotypes were used: C57BL/6 wildtype (WT), and AhR^-/-^^41^, CYP1A1^-/-^^42^, CYP1A2^-/-^^43^ and CYP1B1^-/-^^33^, all backcrossed to C57BL/6. All mice were bred and kept under specific pathogen-free conditions in individually ventilated cages at the central animal facility of Helmholtz Center Munich. Mice were kept in autoclaved, individually ventilated cages (501 cm^2^, IVCs) with stocking density according to the EU guideline 2010/63 with a 12 hour dark/light cycle. Each cage was supplied with autoclaved bedding, bite sticks, nestles and mouse houses. Standard diet (Altromin Spezialfutter GmbH & Co. KG, Lage, Germany) and water were provided *ad libitum*. Cage manipulation took place in laminar flow hoods. Air temperature was 22 ± 2°C and humidity 55 ± 10% with daily control and record. Occasionally, C57BL/6 wildtype animals were bought from Charles River (Sulzfeld, Germany). In this case, mice were allowed to acclimatize for at least one week before sensitization. Both male and female mice over the age of 7 weeks were used.

For the generation of bone marrow chimeras, recipient mice were lethally irradiated by a Co-60 source with two doses of 6 Gray 4 h apart. Irradiated mice were reconstituted with 8 × 10^6^ purified bone marrow cells of respective donors by i.v. injection. After reconstitution, mice received 0.25 mg/ml Enrofloxacin (Baytril; Bayer Vital GmbH) in drinking water for 3 weeks. Mice were maintained for 10-12 weeks to allow replacement of the hematopoietic system before allergen sensitization protocols were applied.

Experiments were performed on age- and sex-matched animals kept in the same rack, according to the European Convention for Animal Care and Use of Laboratory Animals and were approved by local ethics committee and government authorities (ROB-55-2-2532.Vet_02-18-94 and 55.2-1-54-2532-156-12). Human samples were obtained from the BioArchive of the Comprehensive Pneumology Center Munich. All patients gave written informed consent and the study was approved by local ethics committee of the Ludwig-Maximilians University of Munich, Germany (number 19-630).

### Preparation of ragweed and house dust mite extracts

Aqueous ragweed pollen extracts (RWE) were prepared as previously described^44^. Briefly, ragweed pollen were suspended in PBS (2.5 mg/ml), incubated for 30 min at 37°C and then centrifuged for 10 min at 4000 rpm by 4°C. The supernatants were sterile filtered through a 0.22 μm syringe filter (Merck KGaA, Darmstadt, Germany) and frozen at −80°C. House dust mite extracts from *Dermatophagoides farinae* (Stallergene Greer Laboratories, Kamp-Lintfort, Germany and Citeq BV, Groningen, NL) were suspended in 9 % NaCl (1 mg/ml) and stored at −20°C until use. The final solution for intranasal instillation was prepared in PBS.

### Mouse models of allergic airway inflammation

For the ragweed model, a sensitization protocol previously established in BALB/c mice^44^ was adapted to C57BL/6. In short, mice received bilateral intranasal (i.n.) instillations of ragweed pollen extract (RWE) (20 mg/ml; 10 μl/nostril) once a day for a total of 23 days in 5 consecutive weeks. For the HDM model, mice were sensitized once by bilateral i.n. instillations of house dust mite extract of *Dermatophagoides farinae* (HDM) (50 μg/ml; 10 μl/nostril) (Stallergene Greer Laboratories, Kamp-Lintfort, Germany and Citeq BV, Groningen, NL). After eight days, mice were challenged with HDM (500 μg/ml; 10 μl/nostril) for 4 consecutive days. Control animals of both protocols received same amounts of PBS. Weight was monitored throughout the whole experiment. Mice were sacrificed either 24 or 72 h after the last instillation for the RWE or HDM protocol, respectively. After recovery of serum and bronchoalveolar lavage fluid (BALF)^44^, lung tissue was prepared for histology, immunofluorescence, real-time PCR or FACS analysis. Non-lavaged lungs were used for characterization of non-hematopoietic cells by immunomagnetic- or flow cytometry-based cell separation, real-time PCR and metabolite analysis.

### Analysis of plasma immunoglobulins

Plasma samples were prepared from blood taken prior to sensitization (day 0) and at sacrifice (endpoint). Total IgE and RWE-specific IgG1 were measured as previously described^44^. For the detection of Dermatophagoides farinae (Der-f)-specific IgG1, 96-well plated were coated with goat anti-mouse IgG1 (CITEQ Biologics, Groningen, Netherlands) and mouse plasma was added. Subsequently, samples were incubated with biotinylated Der-f (CITEQ), avidin-horseradish peroxidase (Biolegend, San Diego, CA, USA), tetramethylbenzidine (ThermoFisher Scientific, Rockford, IL, USA) and read at 450 nm according to the manufacturer’s instructions.

### Analysis of bronchoalveolar lavage (BAL) and lung histology

BAL and evaluation of inflammatory cell infiltration were performed as described previously^44^. Aliquots of cell-free BAL fluid were used to measure cytokines using a Multiplex-bead-array (Mouse ProcartaPlex, Thermo Fisher Scientific, Waltham, MA, USA) according to manufacturer’s instructions. For lung histology, after BAL, the lungs were excised and the left lobe fixed in 4% buffered formalin and embedded in paraffin. Sections of 4μm thickness were stained with hematoxylin-eosin (H&E) and periodic acid Schiff (PAS). Mucus hypersecretion and inflammatory cell infiltration were graded in a blinded fashion on a scale from 0 to 4 (0=none, 1=mild, 2=moderate, 3=marked, 4=severe), reflecting the degree of the pathological alteration^45^.

### Immunofluorescence

Immunofluorescence staining was performed on mouse and human lung tissue embedded in paraffin. Slides were deparaffinized overnight at 60 °C and rehydrated in a graded series of ethanol and distilled water. For antigen retrieval the slides were immersed in citrate buffer solution (pH 6.0) and placed into a Decloaking Chamber heated up to 125 °C (30 seconds), 90 °C (10 seconds), and cooled down to room temperature. Following a washing step in Tris buffer, blocking with 5% BSA in Tris buffer, the slides were double-stained with CYP1B1 rabbit polyclonal (aa400-450) antibody (LS-B1790, LifeSpan BioSciences, Inc., Seattle, WA, USA) and CC10 mouse monoclonal antibody (E-11) (sc-365992, Santa Cruz Biotechnology, Inc., Dallas, TX, USA). The primary antibodies were diluted in antibody diluent (Zytomed Systems, Berlin, Germany), CYP1B1 (1:50) and CC10 (1:100). For isotype control, rabbit IgG (sc-3888, Santa Cruz Biotechnology, Inc.) and mouse IgG1 kappa isotype control (16-4714-82, Invitrogen, Carlsbad, CA, USA) were used. The staining with the primary antibody occurred overnight in a wet chamber at 4 °C. After rinsing in Tris buffer, a 1:250 dilution of each secondary antibody (Alexa Fluor 488 donkey anti-mouse and Alexa Fluor 568 donkey anti-rabbit, respectively for CYP1B1 and CC10 antibodies, Thermo Fisher Scientific), as well as DAPI (Sigma-Aldrich; St Louis, MO, USA, 1:2000) was applied to the slides and incubated 1 hour at room temperature in the dark. After rinsing again in Tris buffer, the slides were mounted in Fluorescence Mounting Medium (Dako, Hamburg, Germany). Images of the sections were obtained using Axiovert II (Carl Zeiss) and processed using the ZEN 3.2 software (blue edition, Carl Zeiss).

### Flow cytometry of murine lungs

After sacrifice, lungs were perfused with PBS, minced and digested in RPMI 1640 (Gibco, Life Technologies GmbH, Darmstadt, Germany) containing 1 mg/ml collagenase A (Sigma-Aldrich) and 100 μg/ml DNase I (Sigma-Aldrich) at 37 °C for 30 minutes. Digested lungs were meshed and filtered through a 70-μm cell strainer and centrifuged at 500 × g for 10 min. Cell pellets were resuspended in a 40% Percoll (GE Healthcare, South Logan, USA) solution and layered onto an 80% Percoll layer. The Percoll gradient was run at 1500 × g at room temperature for 15 min. The interlayer containing mononuclear cells was collected, washed and incubated with Fc block (BD) and stained with the corresponding antibodies for 30 minutes on ice. Intracellular staining was performed using a Foxp3 fixation/permeabilization kit (eBiosciences, San Diego, CA, USA) according to the manufacturer’s instructions. Live/Dead exclusion was routinely performed using a kit from Life Technologies. Antibodies used for flow cytometry are listed in Supplementary Table S1 and the gating strategy employed for the identification of lung Th2 cells is depicted in Supplementary Fig. S2a.

### Measurement of tryptophane metabolites in lung tissue

Lungs were excised, weighted, cut into small pieces and snap frozen in liquid nitrogen. All solvents used for metabolite analysis were from Sigma Aldrich, LC-MS grade. For the extraction, a mixture of ice-cold methanol:acetonitrile:water (3:2:1, v/v, 600 μl) was added to 100-150 mg frozen tissue. The tissue was lysed using a tissue Lyser (Qiagen), vortex for 1 min at 4–8 °C and incubated for 4 h at −20 °C. After 10 minutes centrifugation at 14,000g at 4–8 °C, the supernatant was collected. Methanol 80% (vol/vol, - 20 °C, 400 μl) was added to the precipitate, each sample was vortexed for 1 minute at 4–8 °C, and incubated for 30 minutes at −20 °C. After centrifugation at 14,000g for 10 minutes at 4–8 °C, the supernatant was collected and combined to the supernatant of the first extraction step from the same sample. After a final centrifugation step at 14,000g for 10 minutes at 4–8 °C, the supernatant was collected and stored at – 80 °C until LC-MS/MS analysis by targeted metabolomics. Before mass spectrometric analysis, 200 μl of the extracts were dried down in a vacuum centrifuge and resolubilized in 100 μl of 5 mM ammonium bicarbonate in water. Reversed phase liquid chromatography-tandem mass spectrometry (LC-MS/MS) was used for the quantification of metabolites by directly injecting 1 ul of the resolubilized mixture onto an Atlantis PREMIER BEH C18 AX VanGuard FIT Column, (2.5 μm, 2.1 x 100 mm; Waters). A RSLC ultimate 3000 (Thermo Fisher Scientific) HPLC system was used, directly coupled to a TSQ Quantiva mass spectrometer (Thermo Fisher Scientific) via electrospray ionization. A 8-minute-long linear gradient was used, beginning with 99% A (5 mM ammonium bicarbonate in water) linearly rising up to 70% B (95% Methanol and 5% 5 mM ammonium bicarbonate in water) employing a flow rate of 100 μl/min. LC-MS/MS was performed by employing the selected reaction monitoring (SRM) mode of the instrument using the transitions (quantifiers) 205.1 *m/z*→ 188.1 *m/z* (tryptophan); 209.1 *m/z*→ 192.1 *m/z* (kynurenine); 190.1 *m/z*→ 144.1 *m/z* (kynurenic acid) in the positive ion mode. Authentic metabolite standards (Merck) were used for determining the optimal collision energies for LC-MS/MS and for validating experimental retention times. Ion chromatograms were interpreted manually using Xcalibur (Thermo Fisher Scientific).

### Immunomagnetic cell isolation of non-hematopoietic cells from murine lungs

For the isolation of CD45− and CD45+ cells lungs were flushed with PBS and digested as described above. CD45− cells were separated by negative selection from CD45+ cells using CD45 MicroBeads and LC separation columns (all Miltenyi Biotech GmbH, Bergisch Gladbach, Germany), according to the manufacturer’s protocol. After controlling for cell purity by flow cytometry, cell pellets were lysed in RNA-lysis buffer (Zymo Research, Freiburg, Germany) and further processed for real-time PCR analysis.

### RNA extraction and real-time PCR

RNA extraction of whole lung pieces or purified cells was performed using the Quick-RNA Miniprep or Quick-RNA Microprep (Zymo Research), respectively, according to the manufacturer’s instructions and RNA was reversed transcribed with a cDNA synthesis kit (Fermentas, Thermo Fisher Scientific). For qPCR, a SYBR green-containing master mix (Roche, Basel, Switzerland) was used. Genes were run in triplicates in a 384well format using a light cycler (Roche) and normalized to two housekeeping genes. The relative expression was calculated with the 2^−ΔCT^ method and typically fold change above control or wildtype are shown as indicated. Primer sequences are listed in Supplementary Table S2.

### Normal human bronchial epithelial cell (NHBE) cell culture

NHBE culture was performed as previously described^38^. Briefly, primary NHBEs (Lonza, Basel, Switzerland) of four genetically independent donors were grown as monolayers in 100% humidity and 5% CO_2_ at 37 °C in serum-free defined growth media (BEGM, Lonza). NHBEs (passage 3) were used at ~80% confluence in 6-well plates. To avoid gene expression changes or influences by growth factors in the BEGM medium, cells were rested in basal medium (BEBM, Lonza) for 12 h, then stimulated with HDM extract at a final concentration of 40μg/mL (CITEQ), recombinant human IL-4 at 50 ng/ml (R&D Systems, Minneapolis, MN, USA) for 6 h. When indicated, cells were pretreated for two h prior to stimulation with the AhR inhibitor CH-223191 (Sigma-Aldrich) at a concentration of 1μM/mL. For RNA analysis, harvested cells were lysed in RLT buffer (Qiagen, Hilden, Germany) containing 1%β-mercaptoethanol (Roth, Karlsruhe, Germany) directly in the cell culture well.

### RNA extraction, quantification and microarray analysis of human NHBE

Total RNA was extracted using RNeasy Mini Kit (Qiagen, Hilden, Germany). RNA quantification and quality assessments were performed by ultraviolet–visible spectrophotometry (Nanodrop Technologies, Wilmington, DE) and the RNA 6000 Nano Chip Kit with the Agilent 2100 Bioanalyzer (Agilent Technologies, Santa Clara, CA, USA). RNA quality of all samples reached a RNA integrity number (Agilent Technologies) of >8.5. Total RNA was amplified and Cy3 labeled by using the one-color Low Input Quick Amp Labeling Kit (Agilent Technologies) according to the manufacturer’s protocol. Hybridization to SurePrint G3 Human Gene Expression 8×60K Microarrays (Agilent Technologies) was performed with the Gene Expression Hybridization Kit (Agilent Technologies).

### Data analysis strategy for microarray

For microarray, data analysis was performed using the Genespring Software GX 14.9.1 (Agilent Technologies) under minimal data reduction constraints (1.5-fold change and P≤0.05 cutoff). Upon data import a standard baseline transformation to the median of all values was performed, including log transformation and computation of fold changes (log2(A/B) = log2(A)-log2(B)). Subsequently, a principle component analysis was conducted and revealed a homogenous component distribution. Compromised array signals (array spot non uniform due to pixel noise of feature exceeding threshold or above saturation) were excluded from further analysis. Genes regulated more than 1.5-fold were further analyzed by using the paired Student’s t-test and filtered for P-value of <0.05). The significantly regulated genes were summarized in entity lists (see Supplementary Table S3). Only differentially expressed probes annotated with a Gene Symbol were considered for further analysis. Probe intensities were averaged in cases where more than one probe per genes was considered differentially expressed, and the lowest p value among these probes was considered for plotting. Genes annotated as protein coding, using the Bioconductor’s AnnotationHub package (Ensemble Human Genome version 99) were used as input for a KEGG map enrichment test as described for RNA-seq data. Microarray data have been deposited in NCBI’s Gene Expression Omnibus (GEO) and are accessible through GEO Series accession number XXX (will be annotated after acceptance).

### Generation of mouse tracheobronchial epithelial Cells (MTEC) cultures

Mouse tracheobronchial epithelial cells (MTEC) were isolated and differentiated as described previously^46^. Briefly, mouse tracheas were surgically removed and placed in 10 ml 0.15% pronase solution (Roche, Basel, CH) and incubated overnight at 4 °C. On the second day, a single cell suspension was generated by gently rocking the digested trachea. To remove fibroblasts, a native selection step was performed by incubating the cell suspension at 37°C in an atmosphere of 95% air, 5% CO_2_ for 5 h in MTEC basic medium containing 10% FCS^46^. Subsequently, supernatants were collected, cells were counted and plated in a 12 well plate (7.5 x 10^4^-1.0 x 10^5^ cells/well) coated with collagen (Thermo Fisher Scientific). Submerged MTEC cultures were incubated at 37 °C in a humidified incubator containing 95% air, 5% CO_2_ for 7-10 days in MTEC proliferation medium containing retinoic acid^46^.

### EROD-Assay

To determine CYP1A1 enzyme activity an EROD Assay was performed. When indicated, cells were stimulated with HDM or the known AhR agonist FICZ (5,11-Dihydroindolo[3,2-b]carbazole-6-carboxaldehyde, Tocris Bioscience, Bristol, UK or Sigma-Aldrich) to activate the AhR and induce CYP1A1 transcription. Prior to the start of the assay, cells seeded in round-bottom cell culture plates (96 well) were washed twice with pre-warmed PBS (37 °C). After aspirating the PBS, the desired volume of EROD reaction mixture [(5 μM Ethoxyresorufin (7-ER, Sigma-Aldrich), 0.5 mM NADPH (Sigma-Aldrich), 1.0 mg/ml BSA (Sigma-Aldrich), 50 mM Tris (pH 7.4)] was pipetted into the plate and subsequently incubated for 20 minutes at 37 °C. To terminate the reaction 2M glycine (Sigma-Aldrich, pH 10.3-10.4) was added in a ~1:1.5 ratio of glycine:EROD mixture volume. Supernatants were spun down at RT for 1-2 minutes to pellet any cellular debris. Subsequently, a fixed fraction of the supernatant was pipetted into a new plate. The plate was read at 535-550nm excitation and 570-590nm emission with a spectrophotometer (InfiniteM200pro, TECAN).

### Isolation of primary mouse alveolar epithelial cells

Mouse primary alveolar epithelial cells were isolated as described previously^47^. Briefly, cardiac perfusion was performed using DPBS (Life Technologies). Subsequently, a cannula was placed in the trachea to infuse, first, dispase (50 U/ml, Roche) for ~45 seconds to allow the enzyme to distribute throughout the lungs without allowing it to spill back out. Next, 1 ml of 1% low-melt agarose (Sigma Aldrich) was gently infused. Lungs were covered with crushed ice for 2 minutes to allow for the agarose to solidify. Individual intact lung lobes were cut away and additionally kept in a dispase solution (50 U/ml, Roche) for 45 min at room temperature on a rocker at 150 rpm. After 45 minutes digested lungs were decanted in complete DMEM/F-12 (Gibco, Life Technologies) with DNase I (Qiagen) for 10 minutes to avoid clumping of cells. Single cell preparations were serially strained though a 100 μm, 70 μm and 40 μm strainers. Primary epithelial cells (singlets, alive, CD45−, EpCAM+ cells) were isolated with a FACSAria III cell sorter (BD) with a cell purity typically around 96%. The gating strategy is depicted in Supplementary Fig. S2b.

### RNA-sequencing analysis of primary lung epithelial cells

Total RNA was extracted from sort-purified lung epithelial cells from wildtype, AhR^-/-^ and CYP1B1^-/-^mice using a RNeasy Micro Kit (Qiagen) and eluted in nuclease-free water. RNA was desiccated to 3 μl and denatured for 3 minutes at 72 °C in the presence of 2.4 mM dNTP (Invitrogen Carlsbad, USA), 240 nM dT-primer* (Metabion, Planegg, Germany) and 4 U RNase Inhibitor (New England Biolabs, Frankfurt, Germany). Reverse transcription and addition of the template switch oligo was performed at 42 °C for 90 min after filling up to 10 μl with RT buffer mix for a final concentration of 1x superscript II buffer (Invitrogen), 1 M betaine (Themo Scientific), 5 mM DTT (Invitrogen), 6 mM MgCl_2_ (Ambion), 1 μM TSO-primer* (Metabion), 9 U RNase Inhibitor (NEB) and 90 U Superscript II (Invitrogen). The reverse transcriptase was inactivated at 70 °C for 15 min and single stranded cDNA was subsequently amplified using Kapa HiFi HotStart Readymix (Roche) at a 1x concentration together with 250 nM UP-primer* (Metabion) under following cycling conditions: initial denaturation at 98 °C for 3 min, 10 cycles [98 °C 20 sec, 67 °C 15 sec, 72 °C 6 min] and final elongation at 72 °C for 5 min. The amplified cDNA was purified using 1x volume of Sera-Mag SpeedBeads (GE Healthcare, South Logan, USA) resuspended in a buffer consisting of 10 mM Tris (Applichem, Darmstadt, Germany), 20 mM EDTA (AppliChem), 18.5 % (w/v) PEG 8000 (Sigma Aldrich) and 2 M sodium chloride solution (Invitrogen). The cDNA quality and concentration was determined with the Fragment Analyzer (Agilent). For library preparation, 1.5 ng cDNA was tagmented using 0.5 μl TruePrep Tagment Enzyme V50 and 1x TruePrep Tagment Buffer L (TruePrep DNA Library Prep Kit V2 for Illumina, Vazyme), followed by an incubation step at 55 °C for 10 min. Next, Illumina indices were added during PCR (72 °C 3 min, 98 °C 30 sec, 12 cycles [98 °C 10 sec, 63 °C 20 sec, 72 °C 1 min], 72 °C 5 min) with 1x concentrated KAPA Hifi HotStart Ready Mix and 300 nM dual indexing primers. After PCR, libraries were purified twice with 1x volume Sera-Mag SpeedBeads (GE Healthcare) and quantified with the Fragment Analyzer (Agilent Technologies), followed by Illumina sequencing on a Nextseq500 with a sample sequencing depth of 30 mio reads on average.

*dT-primer: C6-aminolinker-AAGCAGTGGTATCAACGCAGAGTCGAC TTTTTTTTTTTTTTTTTTTTTTTTTTTTTTVN, where N represents a random base and V any base beside thymidine; TSO-primer: AAGCAGTGGTATCAACGCAGAGTACATrGrGrG, where G stands for ribo-guanosine; UP-primer: AAGCAGTGGTATCAACGCAGAGT

Sequencing reads were pre-processed with bbduk (https://sourceforge.net/projects/bbmap/^48^) to remove adapters still present in the raw reads and for quality trimming. Pre-processed reads were mapped to the mouse genome mm10 using GSNAP version 2020-03-12^49^. RNA sequencing quality was accessed using RNA-SeQC version 2.3.5^50^. Read counts were obtained using featureCounts version 2.0.0^51^. Splice site support and gene annotations were made using the Ensemble mm10 genome release version 99. Gene counts were filtered using the Bioconductor’s AnnotationHub package (R package version 2.18.0) to contain only protein coding genes. The protein coding genes were used for a differential expression analysis using the Bioconductor package edgeR^52^. Genes were considered as differentially expressed when FDR corrected p-values were <= 0.05. Differentially expressed genes were used as input on a gene ontology enrichment analysis using the R package topGO (R package version 2.38.1). Only nodes containing at least 10 genes were considered in the analysis, using the classic algorithm with the Fisher’s exact test, and the p values were corrected using False Discovery Rate (FDR). Additionally, a Fisher’s exact test was used to detect KEGG metabolic maps ^53^that were enriched for differentially expressed genes. This enrichment test was performed in an automated fashion using all available maps, and the resulting p values were also adjusted using FDR correction. Annotations from maps under “Human Disease” and “Drug Discovery” were ignored for the interpretation of the results, and only “Immune System” was used from heading “Organismal systems”. All RNA-seq related calculations were performed using the R environment for statistical computing. The raw data is available at the National Center for Biotechnology Information’s Gene Expression Omnibus database and are accessible through GEO Series accession number GSE158715 (https://www.ncbi.nlm.nih.gov/geo/query/acc.cgi?acc=GSE158715).

### Statistics

Mouse data is displayed as mean±SEM, histological and metabolite analysis as mean±SD. Statistical significance was determined by ANOVA with Tukey’s multiple comparisons or by Student’s unpaired two-tailed t-test with or without Welch’s correction. The analysis was performed by GraphPad Prism software (version 7.0 and 8.0). Statistical analysis of RNAseq and microarray datasets was performed using R language for statistical computing.

## Acknowledgments

The authors wish to thank the animal caretakers of the Helmholtz Center Munich and Benjamin Schnautz, Johanna Grosch and Anela Arifovic for excellent technical assistance. This work was supported by grants from the European Research Council (ERC Starting grant project number 716718 to C.O.) and the Deutsche Forschungsgemeinschaft (grant number OH 282/1-1 within FOR2599 to C.O. and project P07 within SFB1371 to C.O.). R.d.J. was supported by a fellowship from the Humboldt foundation (grant number 1200905 - HFST-P). M.W. was supported by the Christine-Kühne Center (CK-Care). C.H. is supported by a fellowship grant #2019/14245-1, and by grant #2013/07914-8 from the São Paulo Research Foundation (FAPESP). The VBCF Metabolomics Facility is supported by the City of Vienna through the Vienna Business Agency.

## Author contribution

Study design: J.T.B., C.B.S.-W. and C.O. Design and conduction of experiments F.A., R.d.J., M.W., A-M.M., I.F., M.H. and C.O. Data analysis: F.A., R.d.J., M.W., A-M.M. and I.F. Bioinformatic analyses: C.H., U.Z. Supervision: F.A., J.T.B., T.B., J.EvB., C.B.S.-W., C.O. Writing original draft: F.A., R.d.J. and C.O. Review&Editing: All Authors. Funding Acquisition: C.O.

