## Supplementary figures for "Lung epithelial CYP1 activity regulates aryl hydrocarbon receptor dependent allergic airway inflammation"

### Figure S1

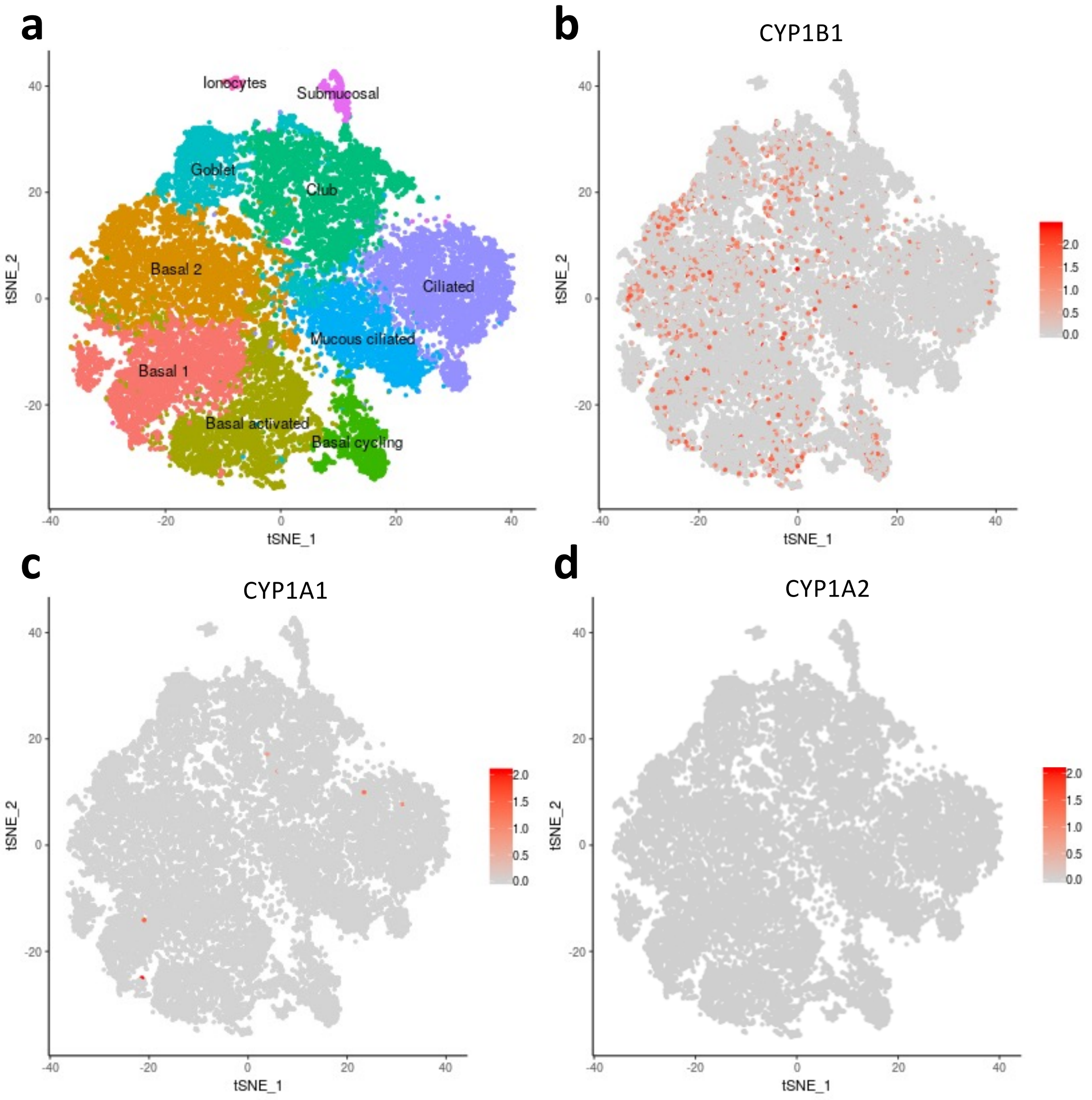


**Figure S1. Expression of CYP1 family members in human asthma airway epithelial cells.**

Data has been retreived from the asthma airway epithelial dataset of the human cell atlas at *https://asthma.cellgeni.sanger.ac.uk*. Map of asthma airway epithelial cell subsets (**a**). Expression of CYP1B1 (**b**), CYP1A1 (**c**), CYP1A2 (**d**) genes among subsets. Colour code for **d)** has been manually changed to grey to better illustrate complete absence of CYP1A2 expression.

Figure S2

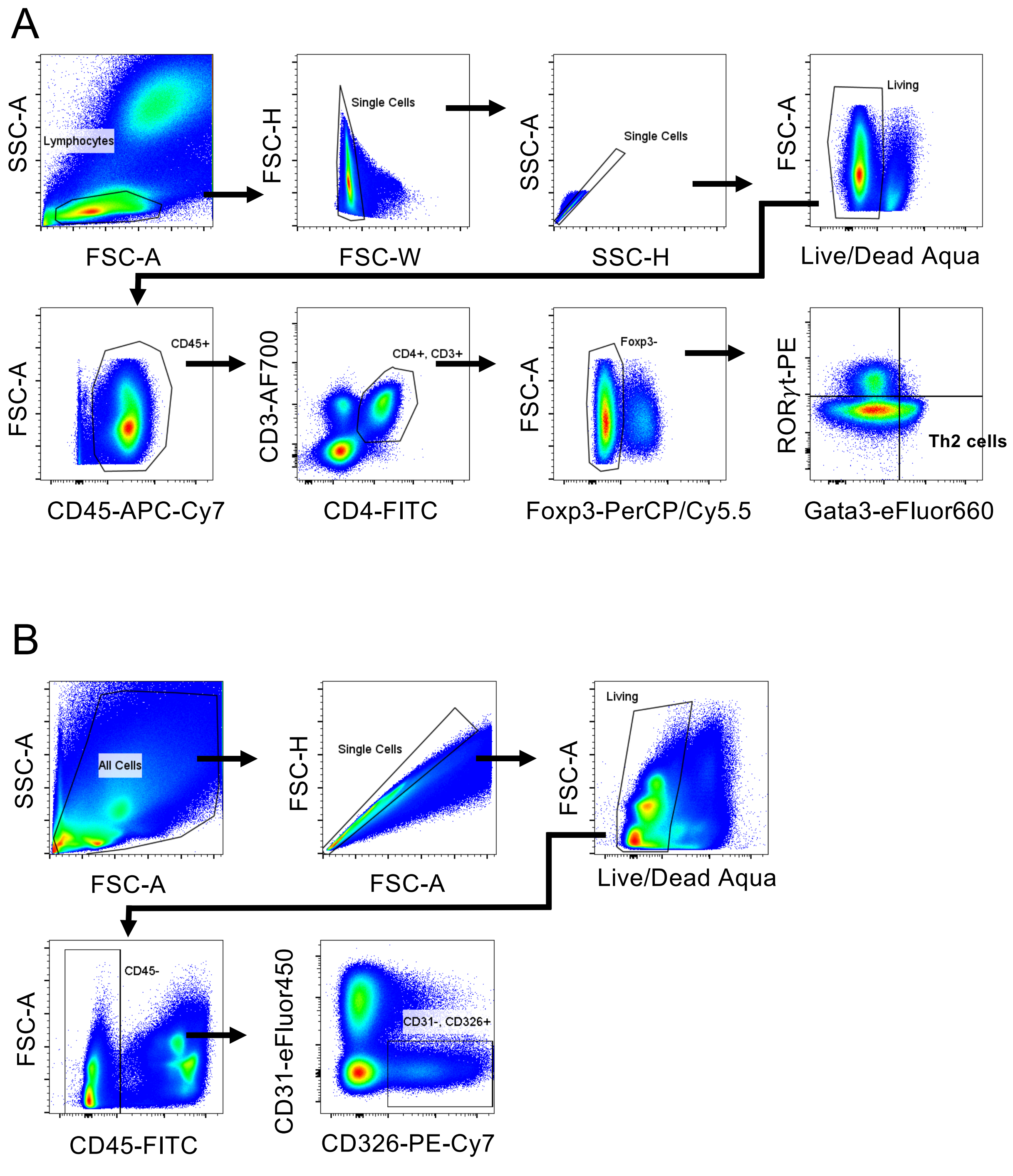


**Figure S2. Gating strategy for the identification of Th2 cells and for the sorting of primary lung epithelial cells.**

Lungs were digested and stained with indicated antibodies as described in material and methods. **A)** Gating strategy for the identification of Foxp3-Gata3+ Th2 cells (related to Fig 1, Fig 2 and Fig 4). **B**) Gating strategy for the flow cytometry-based cell sorting of primary lung epithelial cells of HDM-allergic and control mice (related to Fig 5). Sort purity was routinely above 96%.
